# Attenuated single neuron and network hyperexcitability following microRNA-134 inhibition in mice with drug-resistant temporal lobe epilepsy

**DOI:** 10.1101/2025.06.09.658768

**Authors:** Pablo Quintana-Sarti, Jordan Higgins, Mona Heiland, Amaya Sanz-Rodriguez, Mark O. Cunningham, Omar Mamad, David C. Henshall

**Affiliations:** Department of Physiology & Medical Physics, RCSI University of Medicine & Health Sciences, Dublin, D02 YN77, Ireland; FutureNeuro Research Ireland Centre for Translational Brain Science, RCSI University of Medicine & Health Sciences, Dublin, D02 YN77, Ireland; Discipline of Physiology, School of Medicine, and Future Neuro Research Ireland Centre, Trinity College Dublin, Dublin 2, Ireland

**Keywords:** Anti-epileptic drug, Disease-modification, Epigenetic, Epileptogenesis, Electrophysiology, Patch-Clamp, Noncoding RNA, Hippocampus

## Abstract

The multi-factorial pathophysiology of acquired epilepsies lends itself to a multi-targeting therapeutic approach. MicroRNAs (miRNA) are short noncoding RNAs that individually can negatively regulate dozens of protein-coding transcripts. Previously, we reported that central injection of antisense oligonucleotides targeting microRNA-134 (Ant-134) shortly after status epilepticus potently suppressed the development of recurrent spontaneous seizures in rodent models of temporal lobe epilepsy. The mechanism(s) of these anti-seizure effects remain, however, incompletely understood. Here we show that intracerebroventricular microinjection of Ant-134 in male mice with pre-existing epilepsy caused by intraamygdala kainic acid-induced status epilepticus potently reduces the occurrence of spontaneous seizures. Recordings from ex vivo brain slices collected 2-4 days after Ant-134 injection in epileptic mice, detected a number of electrophysiological phenotypic changes consistent with reduced excitability. Specifically, Ant-134 reduced action potential bursts after current injection in CA1 neurons and reduced miniature excitatory post-synaptic current frequencies in CA1 neurons. Ant-134 also reduced general network excitability, including attenuating pro-excitatory CA1 responses to Schaffer collateral stimulation in hippocampal slices from epileptic mice. Together, the present study demonstrates inhibiting miR-134 reduces single neuron and network hyperexcitability in mice and extends support for this approach to treat drug-resistant epilepsies.

**Significance statement:** Temporal lobe epilepsy is one of the most common forms of drug-resistant epilepsy. Identifying molecular regulators of enduring states of hyperexcitability may lead to new therapeutic approaches. MicroRNAs are short noncoding RNAs that act post-transcriptionally to lower levels of sets of protein-coding genes. Here we show that inhibiting miR-134 reduces spontaneous seizures in mice with active epilepsy. Electrophysiologic recordings from brain slices collected when mice were transitioning to fewer seizures revealed changes to both single neuron and inter-regional communication properties that may explain the reduction in hippocampal network excitability. The findings support the development of this microRNA-targeting approach for epilepsy.

## Introduction

Temporal lobe epilepsy (TLE) is among the most common forms of treatment-resistant epilepsy (Perucca et al., 2023). New therapies are urgently required that work by different mechanisms to traditional anti-seizure medications (ASMs) or that provide long-lasting seizure control (Klein et al., 2024). One of the challenges is that multiple causal pathways may underlie epilepsy and drug-resistance (Loscher et al., 2020). This calls for strategies that enable several pathways to be targeted. RNA-based medicines are an emerging prospect, in particular the targeting of microRNAs (miRNAs) (Morris et al., 2021). MiRNAs serve to fine-tune gene expression by posttranscriptional targeting of protein-coding mRNAs (Henshall, 2024). Following transcription and processing, miRNAs join with Argonaute proteins to form an RNA-induced silencing complex (RISC) before pairing to complementary regions within the 3’ untranslated region of mRNAs resulting in either destabilisation of the mRNA or inhibition of translation (Gebert and MacRae, 2019). Notably, individual miRNAs target dozens and even hundreds of mRNAs, exerting effects across pathways and providing a route to a multi-targeting approach for TLE (Morris et al., 2021)

There is extensive dysregulation of miRNAs and the pathways they control in experimental and human epilepsy (Simonato, 2018; Brennan and Henshall, 2020). This includes miRNAs that regulate inflammation (Aronica et al., 2010; Jimenez-Mateos et al., 2015a; Iori et al., 2017), neuronal morphology (Jimenez-Mateos et al., 2012; Beamer et al., 2018), transcriptional signalling (Tan et al., 2013) and ion channels (Gross et al., 2016), among other processes. Upregulation of miR-134, a brain-specific miRNA originally linked to dendritic spine development (Schratt et al., 2006), has been consistently reported in resected brain tissue from TLE patients, as well as several rodent models (Morris et al., 2019). This includes the mouse intra-amygdala kainic acid (IAKA) model of TLE (Jimenez-Mateos et al., 2012), which has strong translational relevance due to seizures arising from or recruiting the hippocampus, neuropathology, transcriptional overlap with the landscape of human TLE, and drug-resistance profile (Li et al., 2008; Henshall, 2017; Conte et al., 2020; West et al., 2022). Antisense oligonucleotide inhibitors of miR-134, termed antimirs, have been found to reduce evoked seizures in the IAKA model (Jimenez-Mateos et al., 2012), pilocarpine (Jimenez-Mateos et al., 2015b) and pentylenetetrazole (Reschke et al., 2017) models. Importantly, when injected shortly after status epilepticus, antimirs targeting miR-134 potently suppress the later occurrence of spontaneous recurrent seizures (SRS) in rodents (Jimenez-Mateos et al., 2012; Reschke et al., 2017; Gao et al., 2019; Reschke et al., 2021).

The development of a therapy targeting miR-134 for TLE would benefit from advances in two further areas. First, evidence that targeting miR-134 can suppress SRS in animals with pre-existing epilepsy. Here, the IAKA model is well-suited to evaluation since recent studies have demonstrated the SRS are refractory to various ASMs (West et al., 2022; Mamad et al., 2024). Second, additional understanding of the mechanism of action. Previous *in vivo* studies revealed SRS rates increased in antimir-134 (Ant-134)-treated mice co-injected with gapmer oligonucleotides targeting LIM domain kinase 1 (LIMK1) (Reschke et al., 2021). Since LIMK1 regulates the volume of dendritic spines, this indicates that the mechanism may involve changes to local or distant synaptic structures and a re-setting of network excitability. While brain slice recordings have found limited effects of inhibiting miR-134 on neuronal properties in the naïve rodent hippocampus (Morris et al., 2018), it remains unknown whether inhibiting miR-134 alters single neuron or network excitability of the hippocampus in epileptic animals. Here, we show that Ant-134 is potently effective at suppressing SRS in mice with pre-existing TLE and causes a set of electrophysiological changes in the hippocampus that collectively may account for the anti-seizure effects.

## Materials and Methods

### Ethical approval and use of animals

All experimental procedures involving animals were performed according to the European Communities Council Directive (2010/63/EU). Adult male C57BL/6JOlaHsd mice (25–30 g, Harlan) were used under RCSI University of Medicine and Health Sciences’ Research Ethics Committee (REC 1587) approval, and under license from the Ireland Health Products Regulatory Authority (AE19127/P057). Animals were housed on a 12 h light-dark cycle under controlled conditions (temperature: 20°C– 25°C; humidity: 40%–60%). Food and water were available ad libitum.

### Intra-amygdala kainic acid (IAKA) model

The IAKA model was performed as previously described (Mamad et al., 2024). Mice were anesthetised with isoflurane and fixed on a stereotaxic frame. Single-or dual-channel telemetry devices (Data Systems International (DSI), MN, USA) were implanted under the skin in a subcutaneous pocket along the dorsal flank of the mouse. Cables were connected to screws placed on the skull, over the frontal cortex as a reference while the recording electrode(s) was placed over the dorsal hippocampus. A guide cannula was placed on the dura over the right basolateral amygdala (coordinates from adjusted Bregma: anterior-posterior (A/P) = −0.95 mm; lateral (L) = −2.85 mm), for KA injection. Another cannula was placed over the right lateral ventricle from adjusted Bregma: anterior-posterior (A/P) = +0.3 mm; lateral (L) = −0.9 mm), for intra-cerebroventricular (ICV) injections of Ant-134. The antimir (Qiagen) had a phosphorothioate backbone with locked nucleic acid modifications (sequence: TGGTCAACCAGTCACA aligning to nucleotides 1-16 from the 5’ end of miR-134). After 48 h recovery, all animals received an intraamygdala microinjection of KA (0.3 µg in 0.2 µL volume) to trigger status epilepticus. After 40 minutes, mice received intraperitoneal injection of lorazepam ([8 mg/kg]) to reduce mortality and morbidity. IAKA-triggered seizures were confirmed by EEG and mice were then returned to home cages and stayed under climate-controlled conditions and video and EEG monitoring. EEG was recorded from individually housed, freely moving mice for 24 h/day during monitoring. After two weeks baseline recording period, animals received ICV injections of either PBS (epileptic controls) or Ant-134 (1 nmol). Mice were then monitored either for a further two weeks (to record effects on epilepsy) or euthanised 2 – 3 days later and brains processed for electrophysiology recordings (Figure 1).

**Figure 1.**
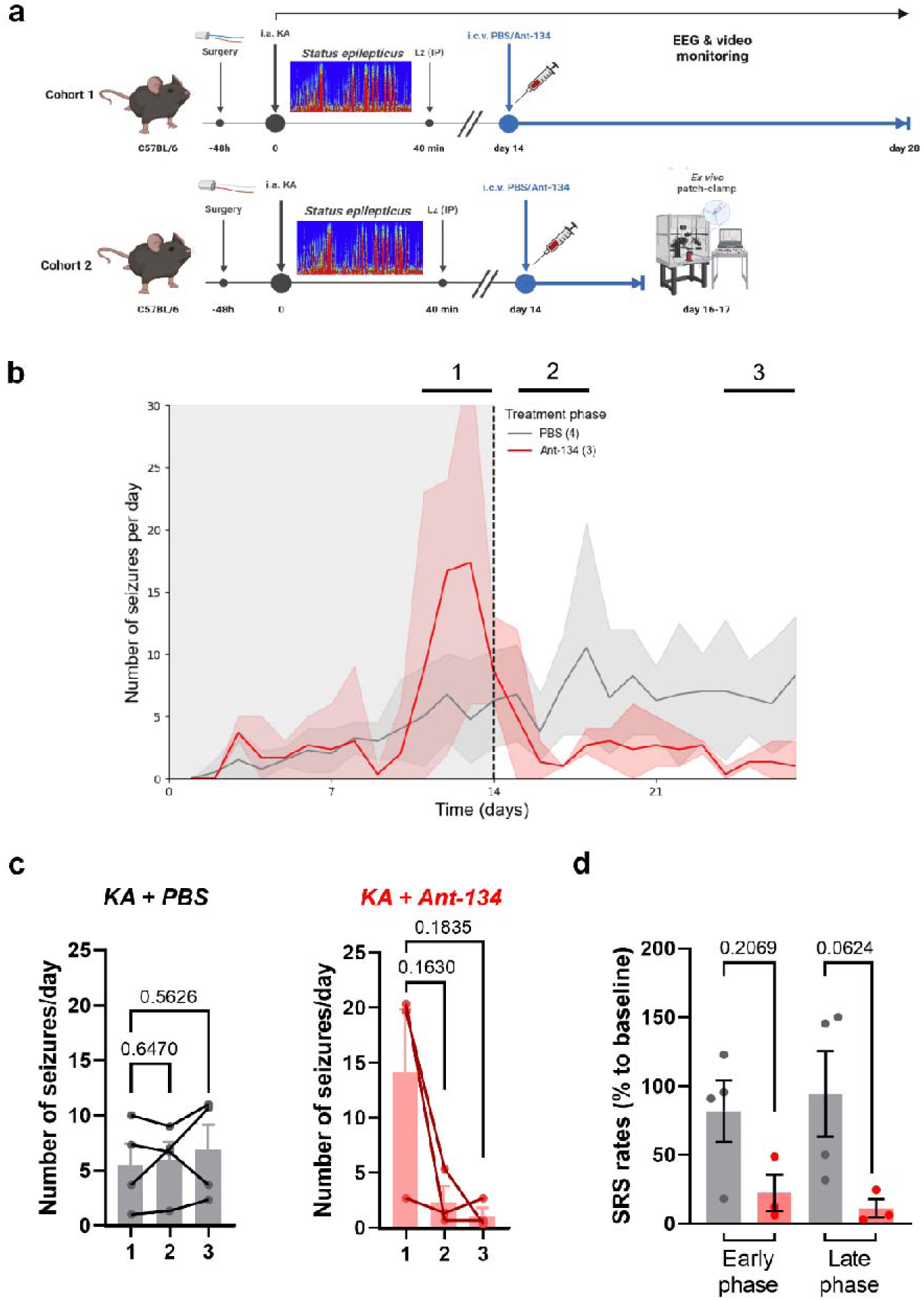
Ant-134 treatment reduces seizures in mice with pre-existing epilepsy. **(a)** Schematic of the longitudinal cohort (top, cohort 1), aimed to study Ant-134 effects on SRS, and the investigative cohort (bottom, cohort 2), aimed to study circuit and cell-specific changes shortly (2-4 days) after Ant-134 administration. **(b)** A single Ant-134 injection (ICV) reduced the number of SRS per day (p=0.089, Two-Way RM ANOVA) relative to PBS-treated epileptic controls during two weeks post-treatment. Three different timepoints were used to further analyse SRS rates: baseline (1), early treatment phase (2) and late treatment phase (3). **(c)** Changes in the number of SRS in individual mice at early (2) and late (3) treatment phases relative to baseline (1) were similar in PBS epileptic controls (left graph) and significantly reduced Ant-134 treated mice (right graph). Paired Student’s *t* tests. **(d)** Drop in SRS rates (% to baseline) at early and late phases of either PBS or Ant-134 treatment. N=3-4 mice/group.

### EEG analysis and counts of SRS

The number and duration of SRS were determined by individually reviewing EEG records, as described previously (Mamad et al., 2024), using LabChart 8 Reader (AD Instruments, Oxford, UK). Spontaneous seizures were defined as high-frequency (>5 Hz), high-amplitude (more than two times baseline) polyspike discharges of ≥10 s duration. Seizure cessation was defined as a return of EEG amplitude and frequency to baseline values with or without postictal amplitude suppression.

### Acute ex vivo mouse slice model

To avoid bias, the treatment administered was unknown at the time of experiment. Briefly, mice were deeply anesthetised (5% isoflurane) and quickly decapitated. The whole brain was subsequently dissected and placed in ice-cold oxygenated (95% O_2_/5% CO_2_) high-sucrose artificial cerebrospinal fluid (aCSF, in mM: 11 Glucose, 1 MgSO_4_, 1 NaH_2_PO_4_, 2,5 KCl, 26.2 NaHCO_3_, 2.5 CaCl_2_ and 119 NaCl, 75 sucrose), and 350 µm coronal slices were obtained with a 7000 smz mkII vibratome (Campden Instruments®, Loughborough, UK). Slices recovered for at least 1 hour in bathing aCSF solution (in mM: 85.6 NaCl, 2.5 KCl, 1.25 NaH_2_PO_4_, 1 MgSO_4_, 2 CaCl_2_, 25 Glucose, 25 NaHCO_3_,100 Taurine, 1600 Thiourea, 3 Myo-inositol, 12.25 N-acetyl-cysteine and 5 Sodium L-Ascorbate, adjusted to 7.2 pH and LJ 280 mOsm) in a recovery chamber at room temperature until recording.

### Whole-cell patch-clamp recordings

After recovery, slices were placed in a submersion chamber in a patch-clamp rig constantly perfused with oxygenated recording solution (in mM: 11 Glucose, 1 MgSO_4_, 1 NaH_2_PO_4_, 2.5 KCl, 26.2 NaHCO_3_, 2.5 CaCl_2_ and 119 NaCl) at physiological temperature (36°C), at a flow rate of LJ 7.5 mL/min. Hippocampal CA1 pyramidal cells were visually identified with an up-right microscope (Scientifica SliceScope) and recorded, digitised at 25 kHz and filtered at 10 kHz or at 2.4 kHz for miniature post-synaptic currents (mPSCs) recordings (Multiclamp 700B, Molecular Devices) (Digidata1440B, Molecular Devices). Borosilicate glass pipettes (5-8 MΩ) were filled with a potassium gluconate based internal solution (in mM: 110 K-gluconate, 40 HEPES, 4 NaCl, 4 MgATP and Na_2_GTP), avoiding cesium-based internal solution to study sodium and A-type potassium currents from the same CA1 neurons. Cells were rejected if resting membrane potential was more depolarised than –55 mV, if series resistance > 30 MΩ, or the series resistance changed by > 20% during the recording. Software automatic series resistance cancellation was performed, and cells rested for at least 5 minutes before performing any experiments. For intrinsic properties, 2-3 neurons per slice were recorded. For the remainder of the experiments, only 1 neuron per slice was used.

### Intrinsic cell properties

Passive membrane properties, including resting membrane potential and cell capacitance, were measured from acquired signals. Action potentials (AP) were induced with a depolarising current steps protocol (–50 to +550 pA in 25 pA increases) and firing frequency and input resistances were measured. Input resistances were directly calculated at each depolarising step following Ohm’s law: *I = V / R*; where *I* was the current injected, *V* the voltage measured and *R* the input resistance. AP properties were measured from threshold APs, which were determined as the first triggered AP at each step, and averaged per recorded cell.

### Miniature EPSCs and IPSCs recordings

Spontaneous and miniature excitatory postsynaptic currents (s/mEPSCs) were recorded in voltage clamp mode while holding the cell at a hyperpolarized holding potential (*V_hold_*) of –80 mV, equivalent to E_GABA_ for 5-10 minutes, excluding all GABAergic inhibitory currents. After sEPSCs recordings, 1 µM tetrodotoxin (TTX) was bath applied to the recording aCSF, to block action potentials and record mEPSCs. Subsequently, with TTX still present in the recording medium, CA1 neurons were held at *V_hold_* +0mV (E_AMPA_) to shift Cl-driving force and enable isolated recording of miniature inhibitory postsynaptic currents (mIPSCs). Neurons were allowed to settle for 5 minutes and mIPSCs were recorded for 5-10 minutes subsequently. Analysis of postsynaptic currents’ inter-event intervals (IEIs) and amplitudes was performed using custom-made event templates in Clampfit software (pClamp, Molecular Devices).

### Input/output curve to electrical stimulation and paired-pulse ratio

To electrically stimulate CA3 inputs to CA1 pyramidal neurons, a bipolar electrode was placed in the CA1 stratum radiatum in the Schaffer collateral (SC) fibres, 200-300 µm lateral to the patch-clamp pipette electrode. Excitatory post-Synaptic potential (EPSPs) were evoked by recording CA1 neurons in current-clamp mode whilst stimulating. Input-output curve was analysed by increasing the stimulation amplitude every 5 sweeps, and averaging EPSPs evoked with the same amplitude. For paired-pulse ratios, two 150 µs electrical pulses were delivered at various inter-stimulus intervals (ISIs) with the cell in current-clamp mode. The probability of neurotransmitter release (NT Pr) was estimated by calculating the ratio between the second and the first evoked EPSPs.

### NMDA/AMPA ratio

In voltage-clamp mode, α-amino-3-hydroxy-5-methyl-4-isoxazolepropionic acid (AMPA)-dependent currents were elicited by electrical stimulation of SC-CA1 pathway by holding the cells at *V_hold_* –70 mV. Subsequently, a mixture of AMPA and N-methyl-D-aspartate (NMDA)-dependent currents was estimated by holding the cells at +40 mV and eliciting currents without modifying electrical stimulation’s amplitude. NMDA-dependent current estimation was measured starting from 50 ms post-stimulation artifact, in efforts to exclude all potential AMPA-dependent current.

### Synaptic plasticity induction and estimation of presynaptic contributions

A theta-burst stimulation (TBS) protocol was used to evaluate the effect of the antimir on the CA3-CA1 circuit plasticity. Briefly, SC fibres were stimulated at basal frequency of 0.2 Hz, until evoking a response of around 3-5 mV of amplitude. After a 10-minute constant baseline response, the TBS was delivered (5 pulses delivered at 100Hz, repeated 20 times at 1 Hz between repetitions). Basal frequency was subsequently returned and recorded for further 30 minutes post-TBS. Averages of EPSP responses within 1 minute post-TBS induction were analysedwithin the last 5 minutes of the 30 minutes post-TBS protocol. Additionally, we performed paired-pulse ratios before the establishment of basal stimulation frequency od EPSPs, and 30 minutes post-TBS. The NT Pr before and after plasticity induction was compared to investigate presynaptic contributions to plasticity between epileptic controls and Ant-134 treated mice.

### Statistical analysis

Statistical analysis was performed with Clampfit v10.7, SigmaPlot v14.0 and GraphPad Prism v8.0 softwares, as appropriate. Two-tailed Student’s t test and One-or Two-Way ANOVA with Bonferroni’s correction for multiple comparisons were used, unless stated otherwise. Repeated measures ANOVA was used as appropriate if there were no missing values, in which case mixed-model effects was applied. Data is presented as mean ± standard error mean. Box and whiskers (min to max) plots are used to illustrate data spread and variability; cumulative probability plots are used to illustrate distribution of large-event experiments and analysed with Kolmogorov-Smirnov cumulative distribution tests. Traces of electrophysiological recordings were obtained with custom software written for this project using Python 3.10 and the pyABF package (https://github.com/swharden/pyABF and see https://pypi.org/project/pyabf).

## Results

### Ant-134 reduces SRS in mice with pre-existing epilepsy

Testing was performed using the IAKA model which is a widely used representation of TLE that displyas partial or full resistance to a range of ASMs including phenytoin, carbamazepine, phenobarbital, valproate, diazepam (West et al., 2022) and levetiracetam (Mamad et al., 2024). We first sought to determine if Ant-134 could reduce SRS in mice with active epilepsy in this model. Mice developed SRS within a few days of IAKA-induced status epilepticus and then, on day 14, received a single ICV injection of Ant-134 (Figure 1a). Rates of SRS dropped within a few days following Ant-134 administration whereas SRS continued in PBS-injected animals (Figure 1b, Table 1). During baseline recordings, epileptic mice in the PBS group experienced 5.5 ± 1.9 SRS/day compared to 14.2 ± 5.8 SRS/day in the Ant-134 before the treatment (Figure 1c). In recordings after injection, rates of SRS were reduced to 1.2 ± 0.7 SRS/day in the Ant-134 group compared to 6.9 ± 2.3 SRS/day in the PBS group, as assessed on day 12 post-ICV injections (Figure 1c). Moreover, at the early (2-4 days) and late (12-14 days) phases after Ant-134 treatment, the SRS rates dropped to ≈ 17% and ≈ 9% compared to pre-treatment SRS rates, respectiveley (Figure 1d, Table 1), while similar rates of SRS continued in the PBS epileptic group (≈ 109% and ≈ 126%, respectively).

**Table 1.**
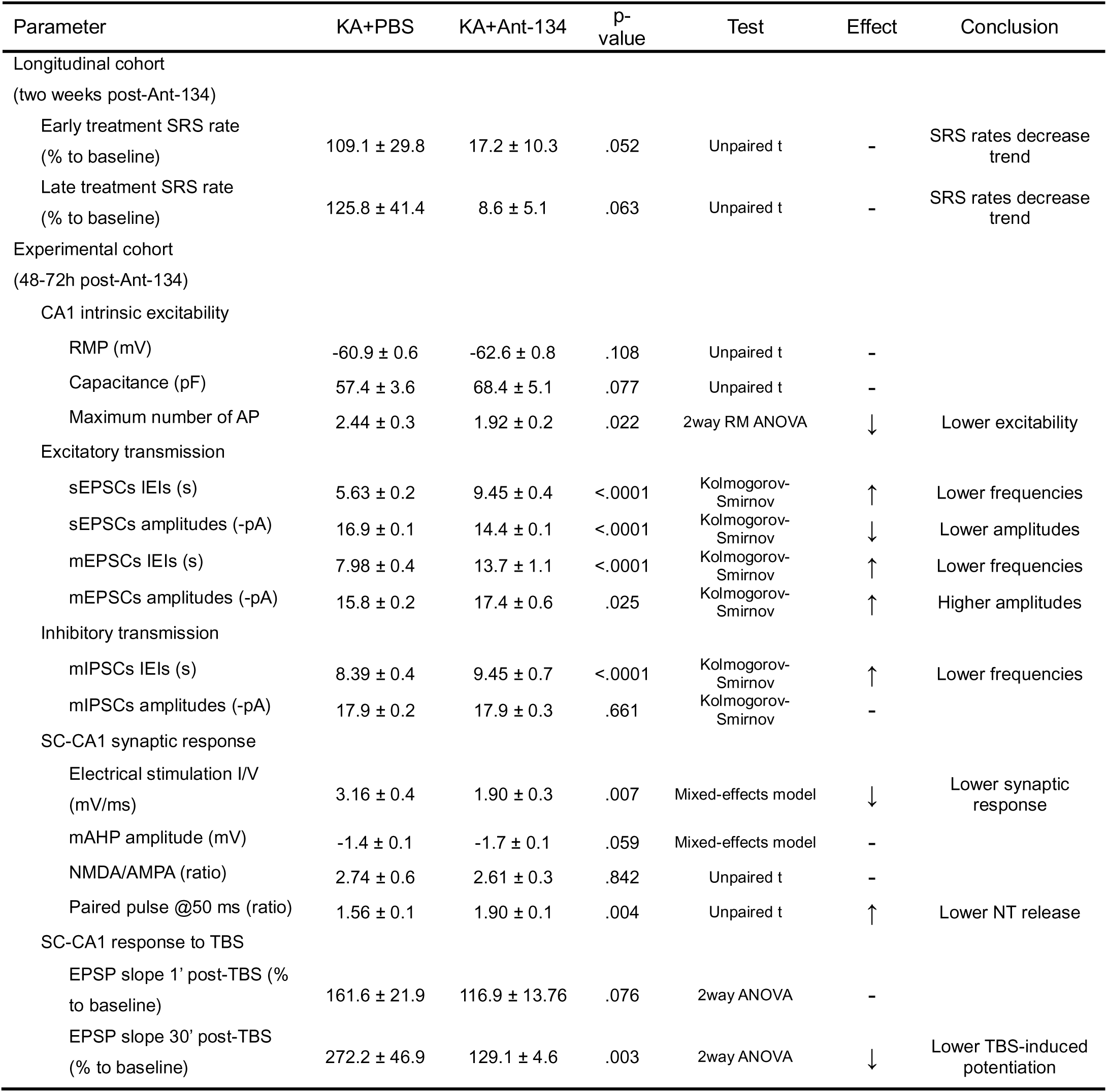
Summary of study’s main findings. Data is presented as mean ± standard error mean.

### Ex vivo brain slice recordings

To explore if Ant-134 changes the properties of the hippocampus in epileptic animals, we performed single neuron and network electrophysiological recordings from acute brain slices. All mice used for electrophysiology studies underwent video-EEG monitoring to confirm the development of epilepsy prior to ICV injections. The appropriate time-point to collect brain slices should take account of the lag from ICV injection to maximal antimir effect while needing to precede when reductions in SRS become significant, the latter of which would make it difficult to separate direct effects of the antimir from secondary effects due to less frequent seizures. We previously reported that knockdown of miR-134 by Ant-134 takes at least 12 h to be significant and peaks 24 – 72 h after injection (Jimenez-Mateos et al., 2012). This coincides with when seizure rates were not yet statistically different between groups (PBS: 6.0 ± 1.6 SRS/day vs. Ant-134: 2.4 ± 1.5 SRS/day; P = 0.181, Figure1b). Accordingly, we selected 2-4 days after Ant-134/PBS injection to collect brain slices for electrophysiological studies. In considering where to direct our electrophysiologic recordings, we avoided the CA3 subfield because this is the main site of neurodegeneration and lesion development in the IAKA model (Mouri et al., 2008; Jimenez-Mateos et al., 2012). In contrast, area CA1 displays only occasional neuron loss in the model while being clinically relevant as a site of neuropathology in human TLE (Blumcke et al., 2017).

### Ant-134 reduces CA1 pyramidal cell burst-firing in epileptic mice

Ant-134 had no significant effect on resting membrane potential (RMP, Figure 2b, Table 1) nor capacitance (Figure 2c, Table 1) of CA1 pyramidal cells in acute brain slices from epileptic mice relative to PBS controls, although we observed a slight trend to higher capacitance values in the Ant-134 group, which could be consistent with an increase in dendritic spine volume (Jimenez-Mateos et al., 2015b). Injection of current into CA1 neurons elicted trains of APs (Figure 2d, e). CA1 neurons from Ant-134 mice exhibited a significantly lower firing rate relative to PBS epileptic control mice, with a ≈ 1.3 fold lower maximum number of action potentials (APs) following depolarizing current injections (Figure 2d,e, Table 1). Input resistances at different current steps were not significantly different in epileptic mice treated with Ant-134 relative to PBS epileptic animals (Figure 2f). These results suggest that miR-134 inhibition reduces intrinsic excitability and likelihood of AP bursting by CA1 pyramidal neurons in epileptic mice.

**Figure 2.**
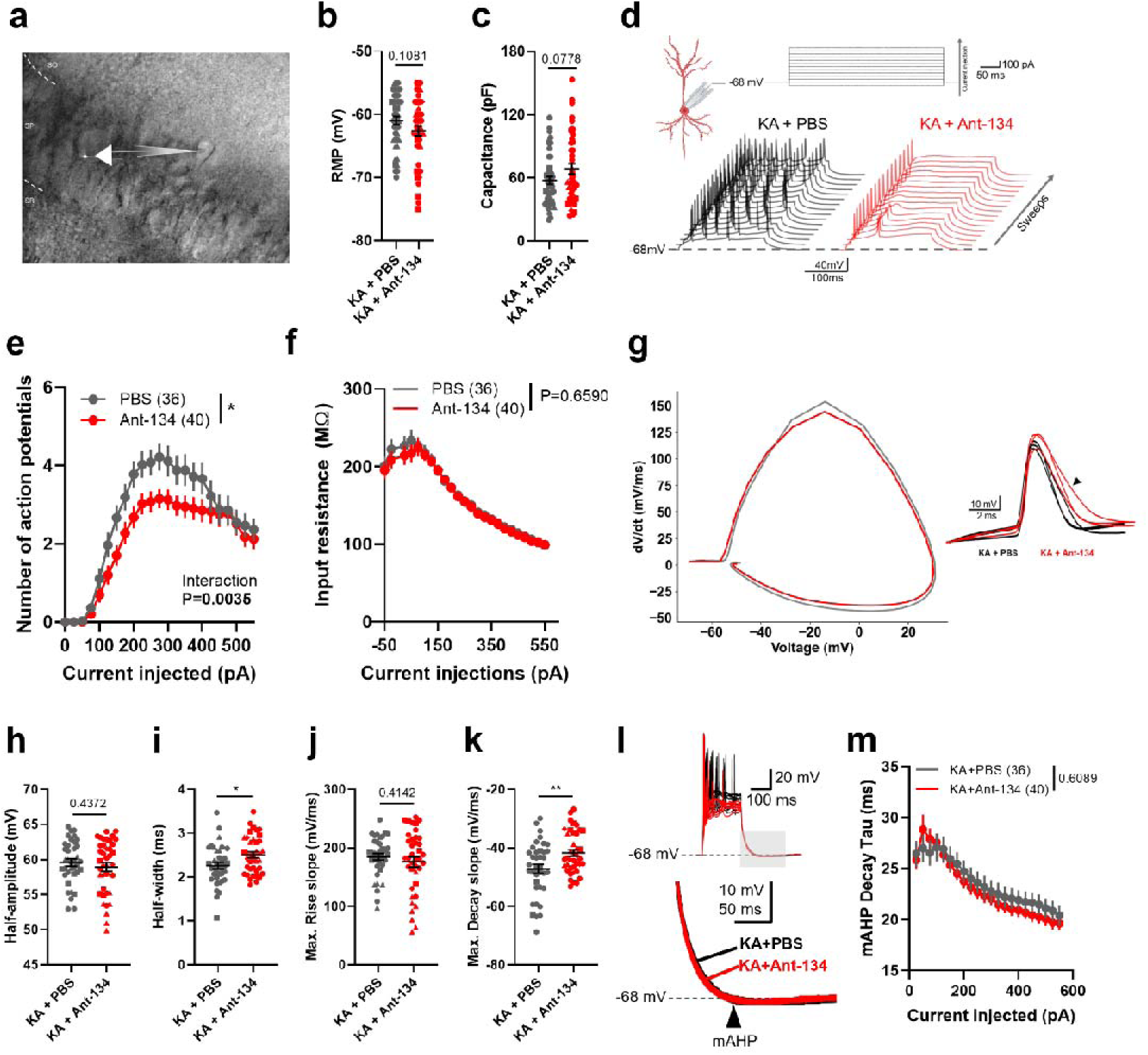
Ant-134 reduces CA1 pyramidal cell intrinsic excitability and modulates action potential properties. **(a)** Representative bright field microscopy image of CA1 area from a brain slice and a patched CA1 pyramidal neuron (electrode, white icon). SO: *stratum oriens;* SP: *stratum pyramidale*; SR: *stratum radiatum*. **(b)** Resting membrane potential (RMP, p=0.11) and **(c)** Capacitance (p=0.07) values of intrinsic properties were similar in both groups (Student’s *t* tests). **(d)** Current-evoked AP firing protocol recorded from CA1 neurons. 200 ms-long square pulses of current were injected between –50 to +550 pA in 25 pA increments to CA1 neurons held at around – 68 mV of membrane potential (top). Representative traces of current-evoked APs at different sweeps (steps) for PBS (dark grey) and Ant-134 (red) CA1 neurons (bottom). **(e)** Input-output curve of current steps protocol showed a reduced (p<0.05, Two-Way RM ANOVA) maximum number of AP firing from CA1 neurons treated with Ant-134. **(f)** Input resistance at each current step was similar in both groups (p=0.66, Two-Way RM ANOVA). **(g)** All-cells average phase-plane plots (left) and traces of 3 representative neurons (right) of current-evoked threshold APs for PBS (dark grey) and Ant-134 (red) treated CA1 neurons. Arrowhead indicates a less pronounced decay slope in Ant-134 neurons. Current-evoked threshold APs exhibited similar half-amplitudes (**h**, p=0.44), longer half-widths (**i**, p<0.05), similar maximum rise slopes (**j**, p=0.41) and less pronounced maximum decay slopes (**k**, p<0.01). **(l)** Representative traces of current-evoked spike mAHPs (top) and zoom-in of the mAHP slope (bottom) from the shaded area. **(m)** Spike mAHP decay *tau* at each current injected was similar between groups (p=0.61, Two-Way RM ANOVA). N=6 mice/group and n=36-40 neurons/group.

### Effect of Ant-134 on other AP properties

Based on the above observations, we further explored AP features in CA1 neurons. The threshold APs, defined as the first triggered AP at each depolarising step, exhibited similar half-amplitude (Figure 2h) and maximum rise slope (Figure 2j) values in epileptic mice treated with Ant-134 relative to PBS controls. Interestingly, CA1 neurons from mice treated with Ant-134 exhibited a significant lengthening of AP duration (Figure 2i) and the decay slope was less pronounced (Figure 2k) in the Ant-134 group relative to PBS controls. We also analysed the medium after-hyperpolarization (mAHP) slope after each spike train following increasing current steps. This showed no difference in mAHP decay *tau* (Figure 2l), peak amplitude, area under the curve, nor time of maximum amplitude (Supplementary Figure S1). Altogether, these results indicate miR-134 knockdown in epileptic mice affects select AP dynamics in a way consistent with reducing CA1 neuron hyperexcitability.

### Ant-134 adjusts excitatory post-synaptic currents

Next, we measured spontaneous excitatory post-synaptic currents (sEPSCs) in CA1 neurons to explore whether incoming excitatory synaptic transmission was altered by Ant-134 treatment (Figure 3a). The average inter-event interval (IEI) between measured sEPSCs in PBS-treated epilpetic mice was 8.8 ± 1.1 s and this was significantly longer in cells from Ant-134-treated epileptic mice (18.3 ± 3.7 s) (Figure 3b, Table 1). Interestingly, we also noted a small reduction in sEPSC amplitude in recordings from Ant-134 mice (14.5 ± 0.5 pA) relative to PBS (16.9 ± 0.9 pA) (Figure 3c, Table 1).

**Figure 3.**
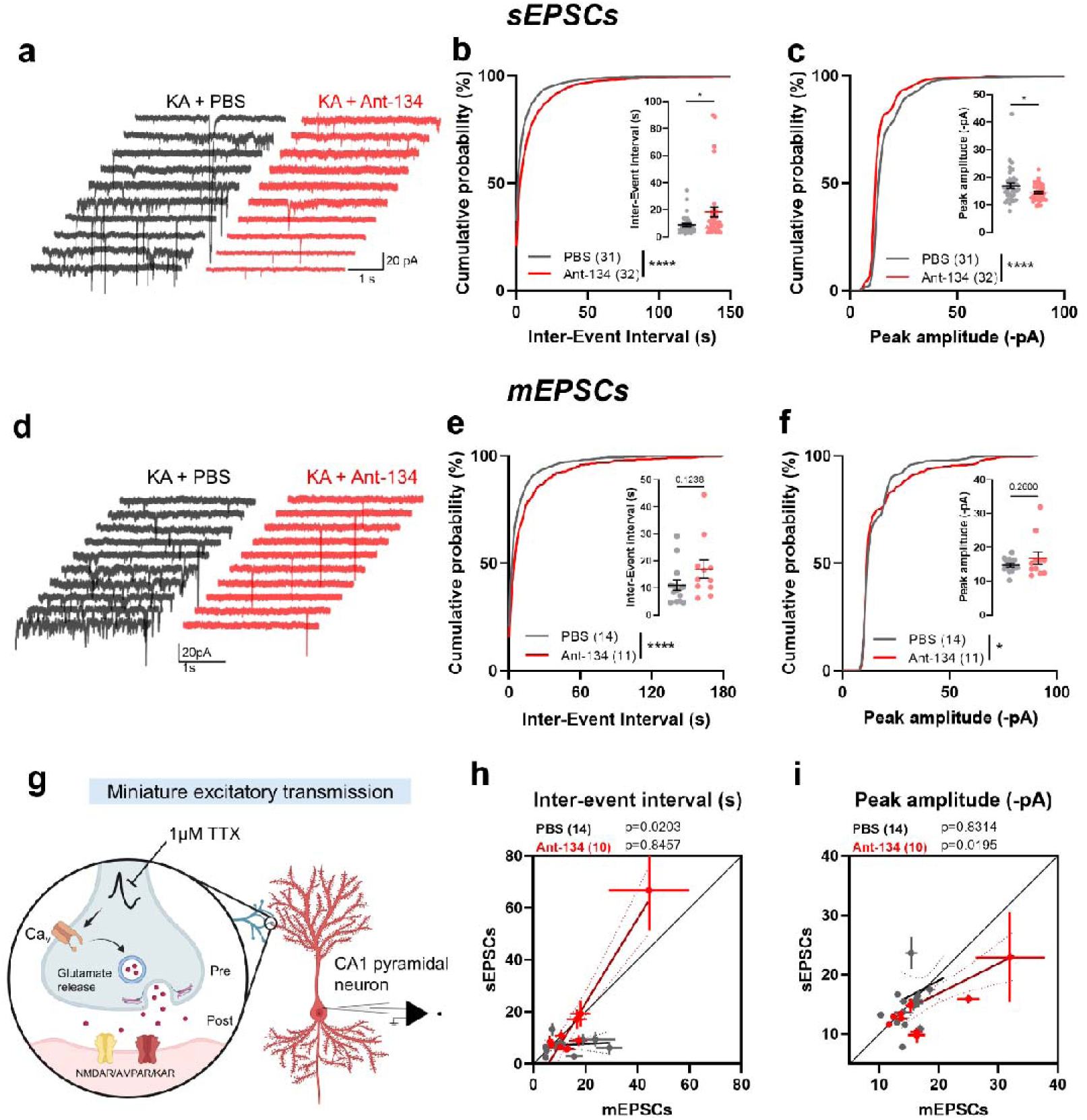
Ant-134 reduces spontaneous and miniature EPSCs in CA1 cells from epileptic mice. **(a)** Representative traces of CA1 neuron spontaneous EPSCs (sEPSCs) recorded from epileptic mice treated with PBS (dark grey) or Ant-134 (red) at a holding potential of –80 mV. Two representative 5-second epochs are shown from one neuron/mouse, 5 neurons/group are shown. **(b)** Cumulative probability of sEPSCs’ inter-event intervals (IEIs) was higher in Ant-134-treated CA1 neurons (p<0.0001, Kolmogorov-Smirnov test). Inset: cell averages of sEPSCs’ IEIs was higher in the Ant-134 group (p<0.05, Student’s *t* test). **(c)** Cumulative sEPSCs’ amplitudes were smaller in Ant-134-treated CA1 neurons (p<0.0001, Kolmogorov-Smirnov test). Inset: cell averages of sEPSCs’ amplitudes were smaller in the Ant-134 group (p<0.05, Student’s *t* test). **(d)** Representative traces of CA1 neurons’ miniature EPSCs (mEPSCs) recorded from epileptic mice treated with PBS (dark grey) or Ant-134 (red). Two representative 5-second epochs are shown from one neuron/mouse, 5 neurons/group are shown. **(e)** Cumulative probability of mEPSCs’ inter-event intervals (IEIs) was higher in Ant-134-treated CA1 neurons (p<0.0001, Kolmogorov-Smirnov test). Inset: cell averages of sEPSCs’ IEIs were similar in both groups (p=0.12, Student’s *t* test). **(f)** Cumulative sEPSCs’ amplitudes were higher in Ant-134-treated CA1 neurons (p<0.05, Kolmogorov-Smirnov test). Inset: cell averages of sEPSCs’ amplitudes were similar in both groups (p=0.26, Student’s *t* test). **(g)** Schema of miniature excitatory transmission onto CA1 neurons. Upon AP occurrence, voltage change in the membrane arrive at the presynaptic voltage-dependent calcium channels (Ca_v_), gating allows calcium influx, vesicle fusion processes and glutamate release to the synaptic cleft. Glutamate binds to postsynaptic glutamate receptors, allowing cation influx into the postsynaptic neuron. Upon tetrodotoxin (TTX) administration, APs are blocked and therefore calcium-dependent release, but quantal, stochastic release occurs. **(h)** Changes in EPSCs’ IEIs before (sEPSCs) and after (mEPSCs) AP termination. EPSCs’ IEIs recorded from PBS controls (dark grey) increased (p<0.05, paired *t* test) after AP termination, not following an identity line (grey line). EPSCs’ IEIs recorded from CA1 neurons from the Ant-134 group (red) do not change (p=0.85, paired *t* test) after AP termination. **(i)** Changes in EPSCs’ amplitudes before (sEPSCs) and after (mEPSCs) AP termination. CA1 neurons’ EPSCs’ amplitudes recorded from PBS controls remained unchanged (p=0.83, paired *t* test) after AP termination, whereas those amplitudes recorded from the Ant-134 group increased (p<0.05, paired *t* test) after AP termination. N=6 mice/group and n=31-32 (sEPSCs) or n=14-11 (mEPSCs) neurons/group.

Changes in sEPSC frequency and amplitude may indicate alterations in pre– and postsynaptic components including neurotransmitter release machinery or the composition of the post-synaptic receptor compartment. To investigate further, we blocked APs using tetrodotoxin (TTX, 1 µM) (Figure 3g). Therefore, the miniature EPSCs (mEPSCs) measured reflect stochastic neurotransmitter release from individual synapses (Figure 3d). The IEI between mEPSCs in CA1 neurons from PBS-treated epileptic mice was 11.0 ± 1.9 s and this was longer in Ant-134-treated samples (IEIs: 16.9 ± 3.7 s) (Figure 3e, Table 1). These results are consistent with a presynaptic change caused by Ant-134. We also observed a shift in the events’ amplitudes before and after TTX administration. The mEPSC amplitudes from CA1 neurons from mice treated with Ant-134 (16.8 ± 1.9 pA) were higher than those in PBS epileptic controls (14.7 ± 0.6 pA) (Figure 3f, Table 1). To further investigate this, we compared both the frequencies and the amplitudes of EPSCs before and after TTX administration in the same cell. After TTX administration, the EPSCs frequencies were reduced in the PBS epileptic animals, as expected. However, those frequencies were not reduced in the Ant-134 group after TTX administration, and remained similar to before AP termination (Figure 3h). Interestingly, amplitudes before and after TTX administration were unmodified in the PBS epileptic group, but increased in the Ant-134 group (Figure 3i). Altogether, our results suggest Ant-134 treatment dampens some excitatory processes in epileptic mice at a single-cell level which may reflect pre– and postsynaptic mechanisms.

### Effects of Ant-134 on inhibitory post-synaptic currents in CA1 neurons

To complement the EPSC findings, we next recorded miniature inhibitory post-synaptic currents (mIPSCs) during patch-clamp recordings from CA1 neurons (Figure 4a). After finishing s/mEPSCs measurements, and with TTX still present in the recording solutions, CA1 neurons were held at *V_hold_*+0 mV to shift the Cl-driving force. Neurons were then left to settle for at least 5 minutes and mIPSCs were measured as outward currents. CA1 neurons from epileptic mice treated with Ant-134 exhibited longer intervals between mIPSCs (12.6 ± 2.7 s) relative to PBS controls (8.1 ± 1.3 s) (Figure 4b, Table 1). The amplitude of mIPSCs were similar between groups (17.5 ± 1.4 pA in Ant-134 vs. 18.6 ± 0.9 in PBS) (Figure 4c, Table 1).

**Figure 4.**
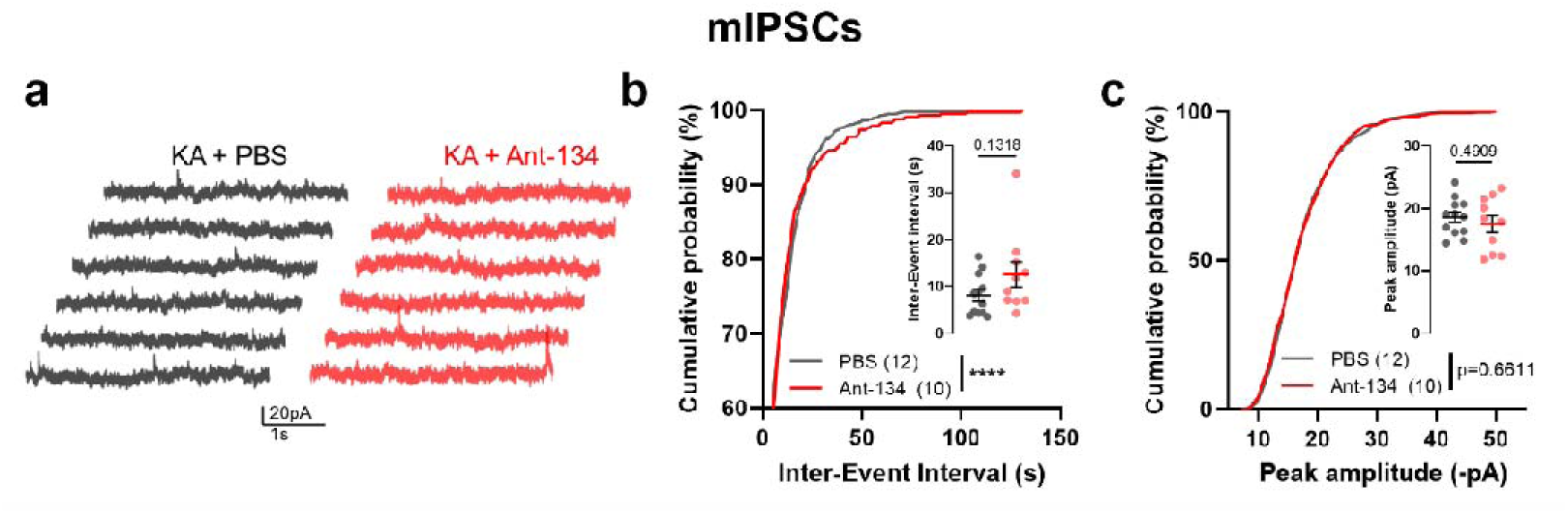
Effect of Ant-134 on miniature IPSCs in CA1 cells from epileptic mice. **(a)** Representative traces of CA1 neuron miniature IPSCs recorded from epileptic mice treated with PBS (dark grey) or Ant-134 (red) at a holding potential of +0 mV. Two representative 5-second epochs are shown from one neuron/mouse, 3 neurons/group are shown **(b)** Cumulative probability of sIPSCs’ inter-event intervals (IEIs) was higher in Ant-134-treated CA1 neurons (p<0.0001, Kolmogorov-Smirnov test). Inset: cell averages of sIPSCs’ IEIs were similar between groups (p=0.13, Student’s *t* test). **(c)** Cumulative sIPSCs’ amplitudes were similar between groups (p=0.66, Kolmogorov-Smirnov test). Inset: cell averages of sIPSCs’ amplitudes were similar between groups (p=0.49, Student’s *t* test). N=6 mice/group and n=12-10 neurons/group.

### Ant-134 effects on Schaffer collateral-to-CA1 synapse stimulation

One of the main excitatory inputs to CA1 neurons is the SC-CA1 synapse, since it completes the cannonical tri-synaptic hippocampal circuit. This synapse is also important for the generation and progression of seizures (Yu et al., 2016). To explore the role of this connection, we perfomed electrical stimulation of SC fibres in slices from epileptic mice while recording EPSPs from CA1 neuronal somata (Figure 5a). We observed a lower stimulus-response curve in SC-CA1 in epileptic mice treated with Ant-134 (1.9 ± 0.3 mV/ms) relative to PBS controls (3.2 ± 0.4 mV/ms) (Figure 5b, Table 1), consistent with reduced general network excitability. Analysis of the repolarization phase of the EPSPs showed no difference in mAHP peak amplitudes between groups (Figure 5c, PBS: –1.4 ± 0.1 mV; Ant-134: –1.7 ± 0.1 mV, Table 1).

**Figure 5.**
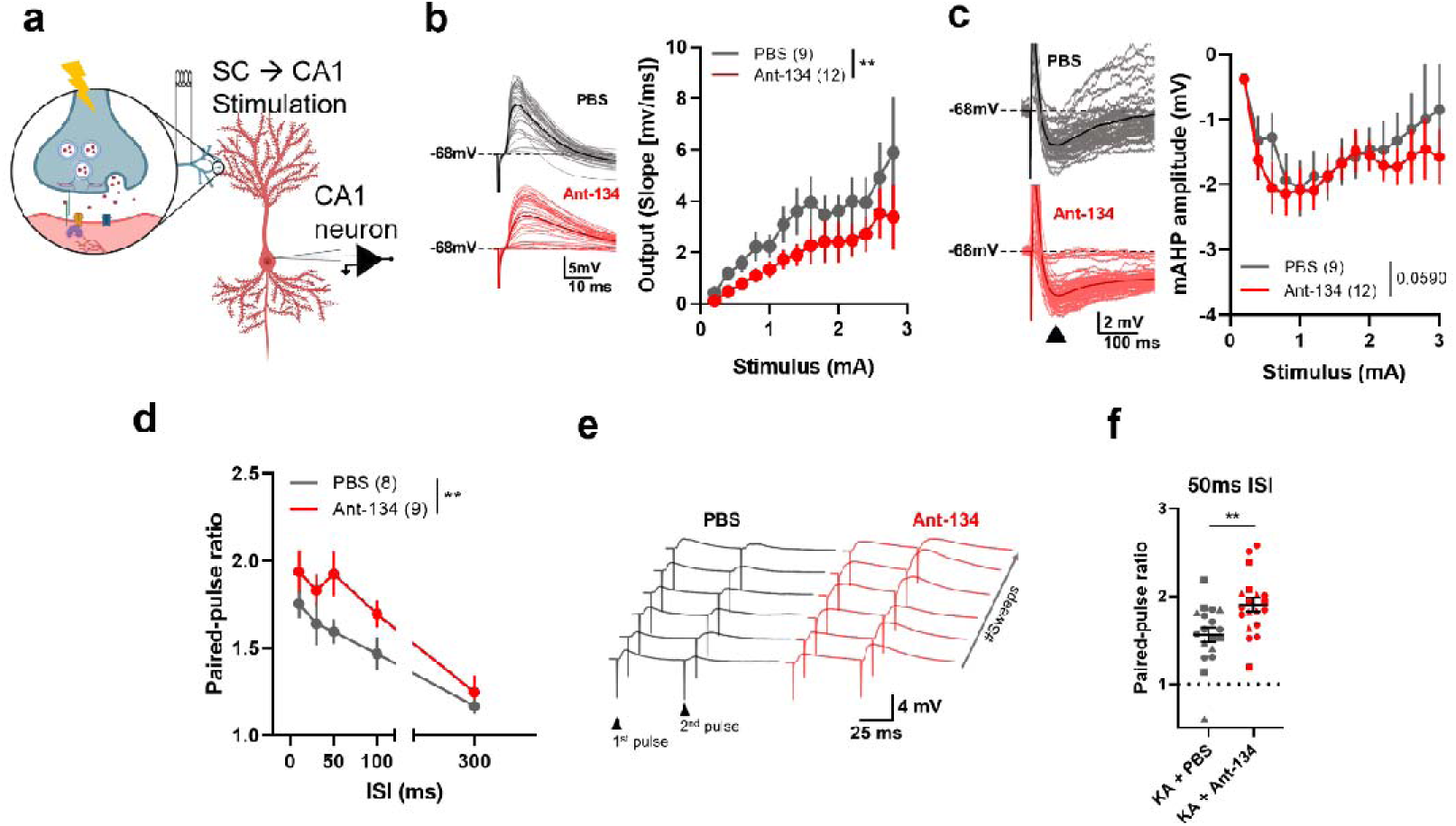
Schaffer collateral-to-CA1 (SC-CA1) synapse response to electrical stimulus in mice treated with Ant-134. **(a)** Schema of experimental protocol. A bipolar electrode was placed in *stratum radiuatum* of CA1 area, stimulating SC fibres and recording the evoked EPSPs in CA1 neurons. **(b)** Representative traces (left) of CA1 neurons’ EPSPs evoked by increasing-amplitude electrical stimulation of SC in both groups. The bold, overlayed traces represent the average trace for each representative cell. Input/output (I/V) curve (right) of EPSP slope following electrical simulation was lower (p<0.01, Mixed-effects test) in the Ant-134 group. **(c)** Representative traces (left) of mAHP observed in the evoked EPSPs’ folllowing electrical stimulation in CA1 neurons from both groups. The bold, overlayed traces represent the average trace for each representative cell. I/V curve (right) of EPSPs’ mAHP amplitude following electrical simulation didn’t change after Ant-134 treatment (p=0.05, Mixed-effects test). **(d)** Electrically evoked SC-CA1 paired EPSPs (paired-pulses) ratios at different inter-stimulus intervals (ISIs) was higher (p<0.01, Two-Way RM ANOVA) in CA1 neurons from epileptic mice treated with Ant-134. **(e)** Traces of paired-pulses at 50 ms ISI recorded from a representative neuron/group. **(f)** Paired-pulse ratios at 50 ms were higher (p<0.01, Student’s *t* test) in Ant-134-treated CA1 neurons. N=6 mice/group and n=9-12 (I/V/mAHP), 8-9 (Paired-pulse ratio) or 18-19 (50 ms paired-pulse ratio) slices/group.

We next stimulated SC-CA1 synapses whilst holding CA1 cells at *V_hold_*–70 mV to measure AMPA receptors (AMPAR)-dependent currents, and sequentially held the cells at +40 mV to measure, after a >5-minutes adaptation period, currents dependent of both NMDAR and AMPAR. We found no significant difference in the NMDAR/AMPAR ratio between groups (PBS: 2.7 ± 0.6 A.U.; Ant-134: 2.6 ± 0.3 A.U.) (Figure S2, Table 1).

### Ant-134 effects on paired pulse stimulation of the SC-CA1 pathway in epileptic mice

We then measured the ratio between two paired electrical pulses (PPR) delivered to the SC at various inter-stimulus intervals (ISIs). This revealed that CA1 neurons from epileptic mice treated with Ant-134 exhibited a significantly longer PPR at different ISIs (1.72 ± 0.127) relative to PBS controls (1.526 ± 0.100) (Figure 5d). We then quantified the PPR at 50 ms in a larger cohort of cells, since it was the ISI at which the PPR difference was greatest. We found a prolongation in the PPR from CA1 neurons treated with Ant-134 (1.904 ± 0.07) relative to PBS controls (1.563 ± 0.08) (Figure 5e,f, Table 1). Altogether, these results suggest that Ant-134 may reduce the SC-CA1 synaptic network excitability in epileptic mice.

### Ant-134 reduces theta-burst stimulation-induced potentiation in epileptic mice

Finally, we examined whether Ant-134 treatment affected synaptic potentiation induced by theta-burst stimulation (TBS) (Grover and Teyler, 1990; O’Connor et al., 1994). TBS mimicks rodents’ in vivo CA1 neurons nested burst firing patterns during resting and spatial learning tasks (Otto et al., 1991). SC fibres were electrically stimulated and the EPSP response was measured in CA1 neurons. After baseline stimulation, TBS was applied and responses were recorded for a further 30 minutes. In epileptic PBS mice, the TBS produced ∼270% increase in EPSP responses relative to baseline. This elevation was attenuated in Ant-134-treated epileptic mice (∼130% relative to baseline) (Figure 6a). TBS-induced short-term potentiation (STP), measured as the EPSP response within 1 minute post-TBS, remained similar in epileptic mice treated with Ant-134 relative to PBS controls (Figure 6b, Table 1), which would suggest that early facilitation processes remain unaltered. The EPSP levels remained lower during the timeline after TBS in recordings from Ant-134 mice relative to controls, and was significantly lower 30 minutes post-TBS (p<0.01, Figure 6c, Table 1). To benchmark the effect of Ant-134 in this TBS protocol, we performed the experiment in brain slices from naïve, non-implanted mice (Figure 6d). The STP component was similar in slices from naïve mice compared to epileptic animals (Figure 6e), suggesting that epilepsy in the IAKA model does not affect the early facilitation processes. Importantly, the EPSP response 30 minutes after TBS induction recorded in naïve animals was similar to Ant-134 treated mice, suggesting Ant-134 may restore normal EPSP potentiation.

**Figure 6.**
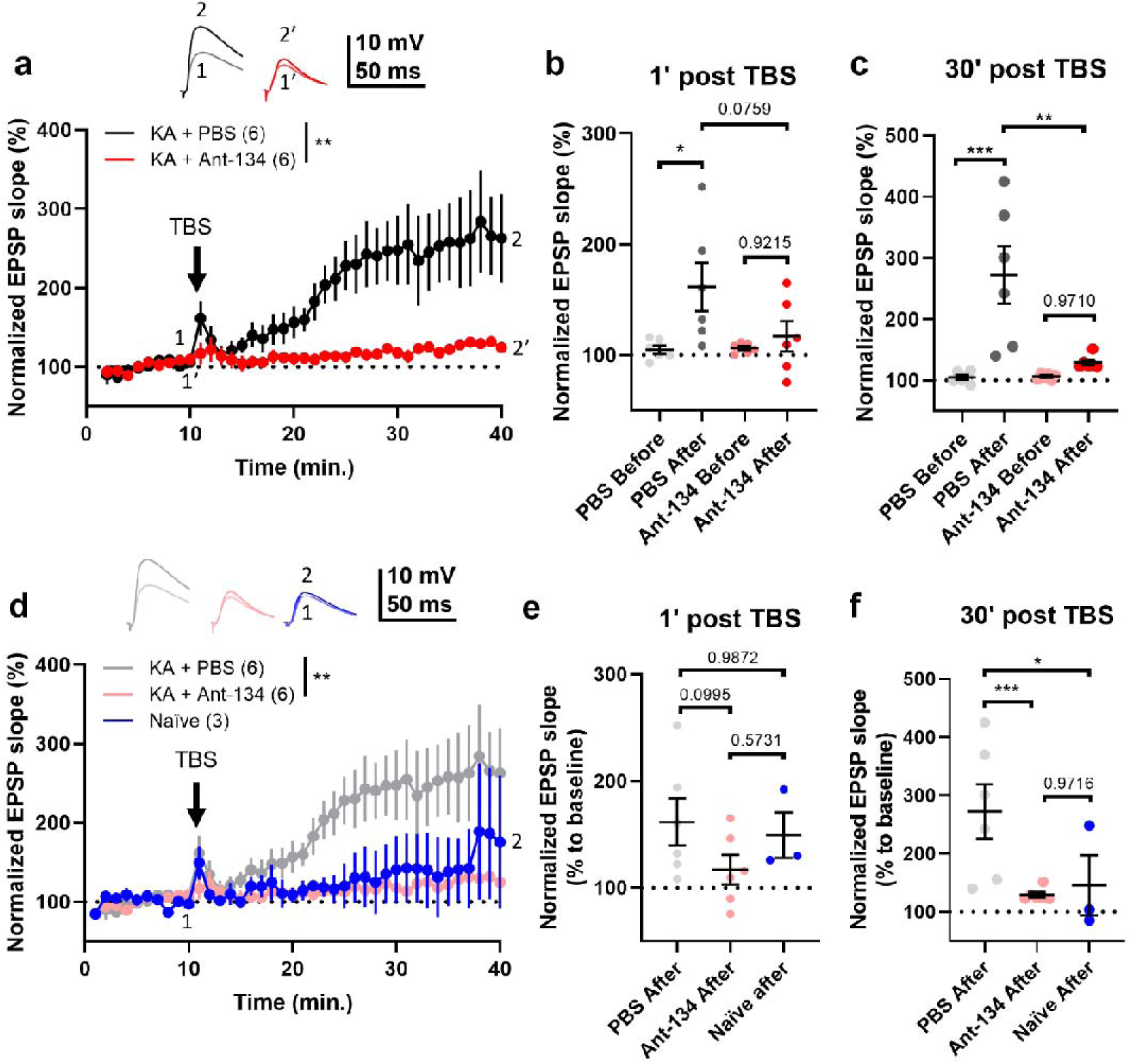
Ant-134 reduces TBS-induced synaptic potentiation. **(a)** Timeline of changes to SC-CA1’s EPSP slope throughout TBS protocol. Top: traces of representative averaged 1-minute response at baseline before TBS (1 for PBS and 1’ for Ant-134) and 30 minutes after TBS (2 for PBS and 2’ for Ant-134). Arrow: TBS induction. EPSP response to TBS was lower in the Ant-134 group (p<0.01, Two-Way RM ANOVA). **(b)** Short-term potentiation (STP) measured at 1 minute post-TBS was similar between groups (p=0.07, pairwise One-Way ANOVA). **(c)** TBS-induced potentiation after 30 minutes was lower in the Ant-134 group (p<0.01, pairwise One-Way ANOVA). **(d)** Timeline of TBS-induction protocol in naïve, non-implanted C57/Bl6 male mice (blue) overlayed to the timeline shown in *(a)*. **(e)** TBS-induced STP in naïve mice was similar to both experimental groups (versus PBS: p=0.99; versus Ant-134: p=0.57, pairwise One-Way ANOVA). **(f)** TBS-induced potentiation after 30 minutes in naïve animals was lower relative to PBS controls (p<0.05, pairwise One-Way ANOVA) and similar to Ant-134 (p=0.97, pairwise One-Way ANOVA). N=3 mice/group and n=3-6 slices/group.

## Discussion

Here, we provide electrophysiological evidence that inhibition of miR-134 reduces SRS in mice with drug-resistant focal epilepsy and produces select changes to single neuron and network excitability properties in the hippocampus from epileptic animals. The study extends the evidence that targeting miR-134 can control seizures, provides insights into the anti-seizure mechanism and supports ‘network therapeutic’ approaches such as miRNA-based treatments as strategies to control hyperexcitable networks in the brain.

The unmet need for novel therapeutic approaches for treatment-resistant epilepsy has driven interest in finding ways to adjust gene expression to restore neuronal network stability (Van Loo et al., 2022; Klein et al., 2024). The targeting of miRNAs enables modest but large-scale adjustment of gene expression within pathways relevant to epilepsy (Brennan and Henshall, 2020; Henshall, 2024). Here, we focus on miR-134, a brain-enriched miRNA originally linked to regulating the development and maturation of neurons and conferring plasticity-related functions via effects on dendritic spines (Morris et al., 2019; Brennan and Henshall, 2020; Soutschek and Schratt, 2023). In both experimental and human TLE, levels of miR-134 are elevated and ASO-based targeting has revealed potent and lasting anti-seizure effects of silencing miR-134 in multiple rodent models (Jimenez-Mateos et al., 2012; Reschke et al., 2017; Reschke et al., 2021). Previous studies, however, tested Ant-134 in either naïve rodents or tracked epilepsy in animals treated immediately after status epilepticus. Thus, it was unknown whether inhibition of miR-134 is effective in animals with active epilepsy.

The first important finding in the present study was that Ant-134 suppressed spontaneous seizures when injected centrally into EEG-monitored male mice previously subject to IAKA-induced status epilepticus, a model which displays secondarily generalised seizures refractory to most classes of ASM (Welzel et al., 2020; West et al., 2022; Mamad et al., 2024). We found rates of spontaneous seizure dramatically declined within 2 – 4 days of a single injection of the antimir. The extent of seizure suppression is similar to results when Ant-134 was given immediately after status epilepticus as an anti-epileptogenic treatment (Jimenez-Mateos et al., 2012; Reschke et al., 2021). Thus, efficacy is not compromised by a need to deliver the treatment before epilepsy is established. This significantly expands the clinical applications. Indeed, there remain major barriers to translating anti-epileptogenic approaches to the clinic because reliable, sensitive and specific biomarkers are not yet available (Simonato et al., 2021; Klein et al., 2025). Ant-134 recently entered large animal testing, with initial results indicating effectiveness in canines with naturally-occurring drug-resistant epilepsy (Henshall et al., 2024). Future studies could test antimirs targeting miR-134 in human models, such as using post-operative human brain tissue (Morris et al., 2022).

Here, we explored how inhibition of miR-134 affects individual neurons and circuits within the hippocampus of epileptic mice. Pathology in the hippocampus of patients with TLE is commonly found within the CA1 subfield (Blumcke et al., 2017), the main function of which is to integrate inputs from CA3 and provide outputs to the subiculum, the latter being the dominant outflow of the circuit (Small et al., 2011). Our brain slice recordings reveal inhibition of miR-134 changes CA1 neuron electrical properties in a direction consistent with lower overall intrinsic excitability. This included attenuated AP firing rates upon depolarising steps, with APs exhibiting slightly longer duration and less pronounced hyperpolarisation phases. Since CA1 pyramidal neuron bursting is a feature of experimental epilepsy and can drive seizures (Marchionni et al., 2019; Mueller et al., 2023), the findings may be relevant to how Ant-134 attenuates SRS in epileptic mice.

Some of the present findings contrast the limited effect of Ant-134 on individual neuron electrophysiological properities in naïve (wild type) rodents (Morris et al., 2018). This suggests that the targets or strength of engagement by miR-134 may change in a disease context. That is, miR-134 may be directed to alternate targets or the uptake of miR-134 or its targets may be altered in the RISC. This is consistent with observations of miR-335 in control and seizure context (Heiland et al., 2023) and from gene disruption studies, where miRNA deletion results in phenotypes mainly in a disease-setting (Bartel, 2018; Lackinger et al., 2019). The target landscape of miR-134 in the brain remains incompletely catalogued but ∼100 different mRNA transcripts bearing 3’ UTR seed sites for MIR134 are present within the RISC from the hippocampus of TLE patients (Heiland et al., 2023). Although the targets of miR-134 have not been mapped in the IAKA model, previous studies indicate miR-134 regulates proteins that support the structure of dendritic spines (Schratt et al., 2006), RNA trafficking (Fiore et al., 2009) and neuronal maturation (Gaughwin et al., 2011). It is likely miR-134 shifts its targeting in the context of epilepsy and engage transcripts encoding proteins that function in neurotransmission, synaptic structure, metabolism and cell-to-cell contact. This aligns broadly with the observed electrophysiologic changes upon inhibition of miR-134 in the IAKA model but suggests other mechanisms may be important. Further studies will be required to identify the specific targets of miR-134 and establish if the rescue of those by Ant-134 contributes to the anti-seizure mechanism(s).

Beyond alterations in the intrinsic excitability of individual neurons, the pathophysiology of TLE features changes to the balance of excitation and inhibition among connections in the hippocampus. Ant-134 had effects on EPSCs, including a reduction in the frequency but slight increase in the amplitude of mEPSCs in recordings from CA1 neurons. These findings are consistent with pre-synaptic alterations of excitatory synaptic transmission in epileptic mice, in addition to the known postsynaptic actions of miR-134 (Schratt et al., 2006; Reschke et al., 2021). Notably, Ant-134 has a small effect to reduce pyramidal spine number (Jimenez-Mateos et al., 2012) but increase spine volume (Jimenez-Mateos et al., 2015b), and anti-seizure effects of Ant-134 can be partially obviated by co-silencing of the miR-134 target LIMK1 (Reschke et al., 2021). Together, this suggests a partial mechanism of Ant-134 is to adjust spines, which would be consistent with the observed trend of higher capacitance values indicating a possible larger neuron surface area. Previous studies of acquired TLE in rodents found increased excitatory synaptic transmission onto CA1 cells (Shao and Dudek, 2004; Clarkson et al., 2020) as well as neurons from the lateral amygdala (Graebenitz et al., 2010). Ant-134 may oppose such pathological excitatory network changes in the IAKA model. We also observed a small reduction in mIPSC frequencies, indicating Ant-134 may adjust inhibitory transmission onto CA1 neurons. Although this seems inconsistent with the potent anti-seizure effects of Ant-134, there is evidence that inhibitory firing rates increase in advance of seizures in drug-resistant epilepsy (Elahian et al., 2018; Miri et al., 2018). Together, the findings indicate Ant-134 may cause subtle shifts in input onto CA1 neurons which may contribute (or be neutral) to its observed potent anti-seizure effects in this model.

The present study identified potential network-level effects of Ant-134 in the epileptic hippocampus. Specifically, the CA1 subfield was refractory to trains of electrical stimuli delivered via the SC input. Ant-134 may be interrupting the passage of high-frequency stimuli, consistent with recordings in non-epileptic rodents (Morris et al., 2018). This would be directly relevant to how Ant-134 reduces SRS in epileptic mice. Notably, kainate models show an increased evoked response in SC-CA1 (Meier et al., 1992; Morin et al., 1998) and increased excitatory neurotransmitter release following hippocampal stimulation is a feature of other epilepsy models (Goussakov et al., 2000; Upreti et al., 2012). Although further studies are requried, these observations, along with the assessments of NMDAR/AMPAR balance and paired-pulse ratio, favor a presynaptic mechanism leading to reduced excitatory neurotransmitter release. The CA1 area has been associated with roles in consolidation and retrieval of long-term episodic memory (Squire et al., 2004; Bartsch et al., 2011) and the TBS protocol findings suggest Ant-134 may aid recovery of synaptic plasticity or cognition which is often impaired in models of epilepsy.

There are potential limitations in the present study. We did not demonstrate which of the various electrophysiological changes were causally important for the effects of Ant-134. Our studies used only male mice and testing should be extended to female animals. We focused our on area CA1 but Ant-134 may produce relevant effects in the CA3 or other subfields. We did not take account of functionally distinct subpopulations of CA1 neurons (Mizuseki et al., 2011; Yao et al., 2023). Ant-134 studies could also focus on neuronal networks beyond the hippocampus. Single cell technologies could enhance insights, for example by identifying the cell type(s) in which targets of miR-134 are de-repressed by the antimir. There may also be opportunities to further optimize antimir formulation, dose and injection schedules.

In conclusion, the present study provides evidence that inhibition of miR-134 potently reduces seizures in a model drug-resistant epilepsy. We also identify changes in the electrophysiologic properties of the hippocampus that may reflect pre-as well as post-synaptic mechanisms. These findings should encourage the investigation of the therapeutic potential for inhibitors of MIR134 to treat drug-resistant seizures in human epilepsy.

## Author contributions

Originial conception of the work: D.C.H. and O.M. Contribution and design of the experiment: P.Q-S., O.M., A.S-R. and D.C.H. Collection of data: P.Q-S, J.H. and O.M. Analysis and interpretation: P.Q-S. and M.H. Drafting the article or revising it for important intellectual content: P.Q-S., D.C.H., M.O.C. and O.M. All authors approved the final version of the manuscript.

## Acknowledgements

The authors thank Dr. Gareth Morris for helpful comments on the manuscript; and Elena Langa, colleagues in the Department of Physiology & Medical Physics, colleagues in the Biomedical Research Facility, and colleagues and the operations team at the FutureNeuro Research Ireland Centre for support.

