## Supplementary Figures for "Attenuated single neuron and network hyperexcitability following microRNA-134 inhibition in mice with drug-resistant temporal lobe epilepsy"

**Supplementary Figure 1.** Ant-134 does not alter spike mAHP parameters in epileptic mice

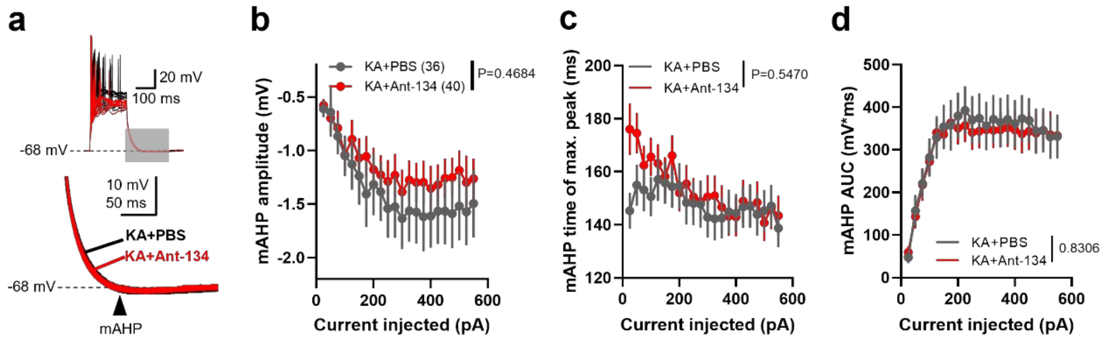


**(a)** Representative traces of current-evoked spike mAHPs (top) and zoom-in of the mAHP slope (bottom) from the shaded area. Spike mAHP amplitude (**b**, p=0.47), time of maximum peak (**c**, p=0.55) and area under the curve (**d**, AUC, p=0.83) remained unaltered following Ant-134 treatment in epileptic mice. Statistics: Two-Way RM ANOVA. N=6 mice/group and n=36-40 neurons/group.

**Supplementary Figure 2.** NMDA/AMPA ratios at the SC-CA1 synapse of epileptic mice treated with Ant-134

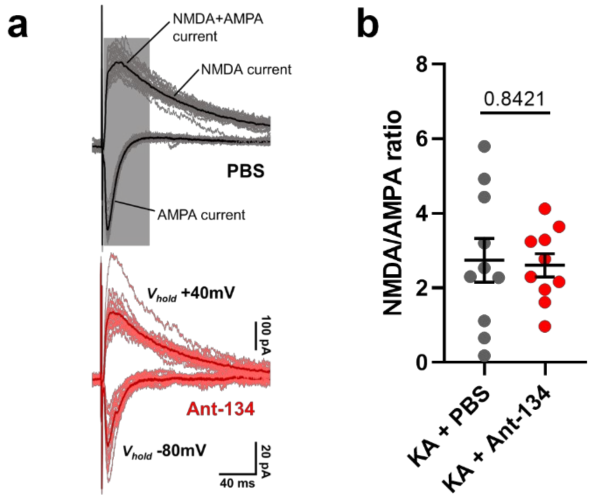


**(a)** Representative traces (left) of electrically evoked SC-CA1 excitatory currents at -80 and +40 mV of $V_{hold}$ to study AMPA- and NMDA+AMPA-dependent currents. **(b)** NMDA/AMPA ratio was similar in both groups (p=0.84, Student's *t* test). N=6 mice/group and n=10 slices/group.
